# Volatile profiling and estimated odor-activity analysis of commercial drug-type cannabis accessions

**DOI:** 10.64898/2026.08.27.747640

**Authors:** Mehdi Babaei, Charles Goulet, Davoud Torkamaneh

## Abstract

Volatile organic compounds (VOCs) define the distinctive aroma of cannabis and critically influence consumer preference, cultivar authentication, and breeding programs. However, systematic characterization of VOC diversity across commercial drug-type cultivars remains limited. This study presents a comprehensive volatilomics-based phenotypic characterization of 165 commercial drug-type cannabis accessions using gas chromatography with flame ionization detection and mass spectrometry (GC-FID/MS). We identified 61 high-confidence VOCs assigned to three biosynthetic classes: terpenoids (*n* = 45), fatty acid-derived volatiles (*n* = 12) and amino acid-derived volatiles (*n* = 4), resolved into 12 subclasses. Analysis of variance revealed highly significant among-accession differences for all compounds (*p* < 0.001; *η*² = 0.67–0.97), with repeatability estimates averaging 0.81 (range 0.50–0.95). Unsupervised clustering partitioned accessions into three distinct chemotypes (*n* = 90, 53, and 22), supported by principal component and t-SNE analyses. Machine learning-based feature selection identified a consensus panel of 12 discriminative compounds (camphene, α-fenchene, sabinene, α-terpinene, (±)-limonene, α- humulene, linalool, endo-fenchol, Δ³-carene, α-thujene, γ-terpinene and α-phellandrene) that recovered the chemotype assignment of 32 of 33 held-out accessions. Estimated odor-activity screening ranked prenylthiol, α-pinene, (±)-limonene, linalool and myrcene highest among the odor-active compounds. All three chemotypes shared a prenylthiol-dominated core (67–77% of summed OAV) and were distinguished by the extent and nature of terpenoid modulation of that core: *minimally modulated* (Cluster ZERO), *citrus–floral modulated* (Cluster ONE) and *pine– terpenic modulated* (Cluster TWO). These findings indicate that volatile diversity in this panel can be summarized by three reproducible chemotypes, providing a quantitative basis for accession characterization and a foundation for future breeding and quality-assessment studies.

## 1. INTRODUCTION

Cannabis (*Cannabis sativa* L.), an annual flowering plant in the Cannabaceae family, ranks among the earliest domesticated crops. Archaeological evidence traces its cultivation to approximately 8000 BCE in East and Central Asia (Babaei et al., 2025, 2022; Crocq, 2020; Ren et al., 2021). Over millennia, humans have used cannabis for fiber, oil, and medicinal applications (Anwar et al., 2006; Barcaccia et al., 2020; Russo et al., 2008; Sun, 2023; Warf, 2014). Twentieth-century global prohibition severely hindered scientific research and breeding progress, limiting our understanding of cannabis biology (Babaei et al., 2026a; Torkamaneh and Jones, 2021; Warf, 2014). Recent legislative shifts have spurred rapid market growth, with the global cannabis market projected to reach US$75.09 billion by 2029 (Statista, 2024).

Cannabis is predominantly a dioecious, diploid species (2n = 20) producing over 545 bioactive secondary metabolites, including more than 120 cannabinoids, over 100 terpenoids, and various flavonoids (Hurgobin et al., 2021; Lapierre et al., 2023b). Legal classification distinguishes hemp (<0.3% Δ⁹-tetrahydrocannabinol (THC)) from drug-type cannabis based on phytocannabinoid content (De Meijer and Hammond, 2005; Sawler et al., 2015). While cannabinoids, particularly THC and cannabidiol (CBD), have drawn primary research focus, volatile terpenoids increasingly shape consumer preferences, therapeutic potential, and product differentiation (Plumb et al., 2022; Russo, 2011). The hypothesized “entourage effect,” where terpenes modulate cannabinoid pharmacology, underscores the importance of characterizing volatile profiles beyond cannabinoid ratios (Booth et al., 2017; Russo, 2011).

Cannabis volatile organic compounds (VOCs) span diverse chemical classes, with terpenoids as the chief aroma contributor (Janta and Vimolmangkang, 2024; Kaminski et al., 2025). Terpenoids derive from two evolutionarily conserved isoprenoid pathways: the plastidial methylerythritol phosphate (MEP) and cytosolic mevalonate (MVA) pathways (Ashour et al., 2010; Oswald et al., 2023; Wink, 2010). Both pathways generate the universal C_5_ precursors isopentenyl diphosphate (IPP) and dimethylallyl diphosphate (DMAPP), which condense to form geranyl diphosphate (GPP, C_10_) for monoterpenes and farnesyl diphosphate (FPP, C_15_) for sesquiterpenes (Semmar, 2024). Terpene synthases (TPS) subsequently catalyze the cyclization, rearrangement, and modification of these linear precursors into structurally diverse terpenoids. Cannabis possesses remarkable biosynthetic capacity, with over 100 distinct terpenes identified to date, governed by at least 30 functional TPS genes from a family of 55 characterized members (Allen et al., 2019; Kaminski et al., 2025). The terpenoid and cannabinoid pathways intersect at GPP, which serves as both the substrate for monoterpene biosynthesis and the prenyl donor for cannabigerolic acid (CBGA) formation, suggesting potential metabolic competition or coordinate regulation (Booth et al., 2017; Lim et al., 2021).

Volatile profiling has emerged as a powerful tool for cannabis chemotype differentiation, with terpenoid profiles providing superior discriminating power compared to cannabinoid ratios alone (Fischedick, 2017; Hazekamp et al., 2016). Studies employing headspace solid-phase microextraction coupled with gas chromatography-mass spectrometry (HS-SPME-GC-MS) have identified distinct patterns across tissues and cultivars (Janta and Vimolmangkang, 2024; Jin et al., 2021). Machine learning (ML) approaches have demonstrated utility for cannabis trait prediction (Babaei et al., 2026b; Yoosefzadeh Najafabadi and Torkamaneh, 2025), yet its systematic use for volatile chemotype classification remains limited.

Beyond abundance, volatile sensory impact is determined by odor activity value (OAV), calculated as the ratio of compound concentration to its odor detection threshold (ODT) (Patton and Josephson, 1957; Ruth, 1986). OAV provides a quantitative framework bridging analytical chemistry and human olfactory perception, widely applied in flavor chemistry of wine, beer, and other aromatic matrices (Pu et al., 2025). Applied to cannabis via multidimensional GC-MS- olfactometry, this showed low-concentration volatiles (e.g., certain aldehydes, thiols) exert disproportionate influence, shifting focus from major terpenes like myrcene and limonene (Rice and Koziel, 2015).

Despite the importance of volatiles in consumer perception and product quality, knowledge gaps persist in breeding and quality control programs. Systematic, quantitative characterisation of volatile diversity across large germplasm panels remains scarce, and objective chemotype frameworks based on absolute volatile concentrations are lacking. This study bridges these gaps via comprehensive volatilomics phenotyping of 165 commercial drug-type cannabis accessions, integrating quantitative volatile profiling, chemotype classification, and odor activity value (OAV) analysis to establish a framework for cultivar characterization, authentication, and breeding.

## 2. MATERIALS AND METHODS

### 2.1 Plant materials

All research activities, including procurement and cultivation of *Cannabis sativa* L. plants, were conducted under Health Canada research license (LIC-QX0ZJC7SIP-2021) in full compliance with federal regulations. The study utilized a panel of 165 drug-type cannabis accessions previously characterized by Lapierre et al. (2023a). Plants were propagated and grown in high-performance greenhouse facilities at Université Laval (Québec, QC, Canada) under standardized commercial production conditions (Lapierre et al., 2023a). Female inflorescences were harvested at physiological maturity (9-10 weeks post-flowering induction) and immediately subjected to volatile profiling.

### 2.2 Volatile organic compound (VOC) extraction and chemical analysis

#### 2.2.1 Volatile profiling by GC-FID/MS

For each accession, 4–10 g of fresh female inflorescences was harvested and coarsely trimmed. From this pool, 7–10 fragments were combined to give a representative subsample of 1 g, which was placed in a sealed extraction tube. Independent subsamples were analyzed per accession as technical replicates (one to five per accession; mean = 3), giving 490 analytical runs in total. Volatiles emitted from each subsample were collected on an adsorbent column and eluted with 0.15 mL of methylene chloride containing nonyl acetate as an internal standard (1.08 µg per gram of sample).

Gas chromatography-flame ionization detection (GC-FID; Agilent Technologies 7890B, Santa Clara, CA, USA) was performed on a DB-5 column (30 m × 0.25 mm i.d. × 0.25 µm film thickness; product no. 122-5033, Agilent Technologies) using hydrogen as carrier gas at constant flow. The temperature program was: 35 °C (hold 1 min), increased to 250 °C at 6 °C min^-1^ (hold 2 min; total run time 38.83 min). The FID temperature was 280 °C.

Compound identification was complemented by gas chromatography-mass spectrometry (Agilent 7890B coupled with 5977B MSD) using a DB-5MS UI column (30 m × 0.25 mm i.d. × 0.25 µm film thickness; product no. 122-5533, Agilent Technologies) and helium carrier gas at 1.2 mL min^-1^. The oven temperature program was: 35 °C (hold 1.46 min), increased to 250 °C at 3 °C min^-1^ (hold 3 min; total run time 76.13 min). Mass spectra were acquired in electron ionization mode at 70 eV, with high-efficiency source temperature 230 °C, quadrupole 150 °C, scan range 35–550 m/z at 1.562 u s^-1^, and solvent delay of 3 min.

#### 2.2.2 Compound Identification

Compounds were identified from their mass spectral data acquired on a subset of reference samples, with reference to commercial mass spectral databases (Adams, 2007; Joulain and König, 1998; Mondello, 2015; Tkachyov, 2008), and based on their expected elution order on a non-polar 5% phenyl/95% methylpolysiloxane capillary column for cannabis (St-Gelais et al., 2024). Peaks were integrated manually by superimposition of all acquired chromatograms using Unichrom 5.0.19.1146, with macros to propagate peak integration across all samples and manual review of the resulting integrations to account for any sample-specific behavior.

Four peaks corresponded to co-eluting compounds and were reported under the dominant constituent: Δ³-carene (with hexyl acetate), limonene (with β-phellandrene and 1,8-cineole), terpinolene (with p-cymenene and fenchone), and selina-4(15),7(11)-diene (with (E)-α- bisabolene). In accessions AO-58 and AN-44, linalool and terpinolene were entirely overlapped, and the same behaviour was observed in one replicate each of AJ-19, AJ-44 and AN-91; the affected peaks were summed for these accessions. Two compounds were assigned as sesquiterpenes from their mass spectra but could not be structurally resolved; they are reported under their internal codes CASA II and BOCA IV. The final dataset comprised 61 compounds quantified across all 165 accessions.

#### 2.2.3 Quantification approaches and dataset selection

Four quantification datasets were derived from the same chromatographic data (internal-standard-normalized, Table S1; external-calibration, Table S2; internal-calibration, Table S3; spike-corrected, Table S4). In the first, peak areas were normalized to the internal standard without calibration. In the second, compounds were quantified in equivalents of nonyl acetate using a nine-point external calibration curve (0.7–54 µg mL^-1^, R^2^ = 0.9977). In the third, compounds were quantified in equivalents of nonyl acetate assuming a linear detector response, using nonyl acetate as an internal standard spiked at 6.97 ng µL^-1^ (1.08 µg per gram of sample). In the fourth, the external calibration was applied, and an injection-specific correction factor was then derived by comparing the calculated concentration of nonyl acetate with its known spiked amount and was applied to all other compounds. Concentrations in the latter three datasets are expressed as estimated concentrations in µg g^-1^ of fresh inflorescence.

The internal-standard-normalized dataset had no missing values, as low-signal peaks had been integrated at baseline; upon conversion to concentration, however, these baseline values became non-physical for a subset of compounds with low response factors in the three calibration-based datasets, and were therefore treated as non-detects (Table S5).

The four datasets were compared for reproducibility and for agreement in accession ranking (Table S5). Within-accession coefficients of variation, reflecting technical reproducibility, were comparable across datasets (median 12.3–17%). Between-accession coefficients of variation, reflecting biological dispersion, were 105%, 85%, 93.7% and 81.6% for the internal-standard-normalized, external-calibration, internal-calibration and spike-corrected datasets, respectively. Spearman rank correlation of accession values was near-unity among the internal-standard-normalized, internal-calibration and spike-corrected datasets (mean *ρ* = 0.967–0.997), whereas the external-calibration dataset showed a distinct ranking pattern (mean *ρ* = 0.883–0.944 against the others).

Variability in the nonyl acetate peak area across all 490 injections was substantial (CV = 37.6%; Table S5), attributed by the analytical laboratory primarily to procedural factors during sample preparation rather than to a fully characterized source (Section 2.2.1). The three datasets that agree closely with one another are all computed from this peak area, so their mutual agreement reflects a shared input rather than independent confirmation. The external-calibration dataset, which does not depend on this peak area, was retained for all subsequent analyses. All four datasets nonetheless yielded the same optimal cluster number (*K* = 3)

#### 2.2.4 Compound classification

Chemical structures and physicochemical properties (molecular weight, molecular formula, octanol-water partition coefficient (LogP), topological polar surface area (TPSA), hydrogen bond donor/acceptor counts, number of rotatable bonds, and canonical SMILES notation) were retrieved from the PubChem database (https://pubchem.ncbi.nlm.nih.gov/) using CAS Registry Numbers and PubChem CIDs. Compounds were assigned to three main classes according to their biosynthetic precursor — amino acid-derived volatiles, fatty acid-derived volatiles, and terpenoids — and further resolved into 12 subclasses on the basis of their structural features and precursor: isoleucine-derived and leucine-derived (amino acid-derived); lipoxygenase-derived and acyl-CoA esters (fatty acid-derived); and acyclic, monocyclic and bicyclic monoterpenes, oxygenated monoterpenoids, sesquiterpenes, uncharacterized sesquiterpenes, acyclic terpenoid thiols and acyclic terpenoid esters (terpenoids) (Ashour et al., 2010; Lange and Turner, 2013; Lei et al., 2021; Wink, 2010). Complete chemical annotations are provided in Table S6.

### 2.3 Odor Activity Value (OAV) calculation

Air-based odor detection thresholds (ODTs) were compiled from *Leibniz-LSB@TUM Odorant Database* (https://www.leibniz-lsb.de/) and *VCF Online Flavor Database* (https://www.vcf-online.nl/VcfHome.cfm), prioritizing values determined by triangle odor bag method (mg/m³; Association 1989; Nagata and Takeuchi 2003). Compound identities were matched using CAS Registry Numbers. For 33 volatile compounds with available ODT data, odor activity values (OAV) were calculated as the ratio of estimated concentration to odor detection threshold, following by Rice and Koziel (2015):

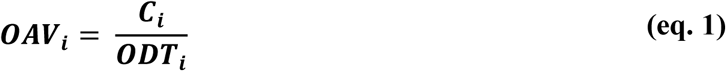

where:

*OAV_i_* = Odor activity value for compound *i*
*C_i_* = Concentration for compound *i*
*ODT_i_* = Odor detection threshold in air for compound *i*

Because Cᵢ is expressed per gram of fresh inflorescence while ODTᵢ is an air-based threshold, this ratio is not dimensionally equivalent to a conventional OAV and does not account for headspace partitioning. It is used here as a relative index to rank compounds and compare chemotypes on a common basis, rather than as an estimate of absolute sensory potency.

OAV values were computed from estimated concentrations (accession means of the available replicates), with non-detects retained as missing. Full details, including ODT values, CAS numbers, chemical categories, sensory descriptors, and original references, are provided in Table S7. Substantial inter-individual variability in ODTs (coefficient of variation 30-100%; Cain and Gent 1991) is acknowledged.

Differences in OAV among chemotypes were tested for each compound by the Kruskal–Wallis test. Compounds characteristic of a given chemotype were identified by one-versus-rest Mann–Whitney U tests with Benjamini–Hochberg correction (Benjamini and Hochberg, 1995), using Cliff’s delta as the effect size (Cliff, 1993). Chemotype profiles were compared using two complementary representations: log-transformed OAV without scaling, and log-transformed OAV Pareto-scaled ((x-μ)/√SD) across all 165 accessions (Van den Berg et al., 2006). The relative contribution of each odor category to the summed OAV of a chemotype was calculated to describe its compositional balance.

### 2.4 Statistical analyses

#### 2.4.1 Dataset Preparation

All statistical analyses were performed on the external-calibration dataset (µg g⁻¹). Best linear unbiased predictors (BLUPs) for each volatile compound were estimated from its technical replicates using the following mixed linear model:

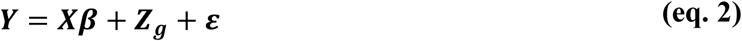

where ***Y*** is the vector of measured concentrations, ***β*** corresponds to the fixed intercept representing the overall mean, and ***g*** corresponds to the vector of random accession effects, assumed to follow a normal distribution with mean zero and variance 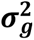 . The ***ε*** accounts for unexplained variation and is modeled as normally distributed with variance 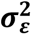, capturing analytical variation among technical replicates. The incidence of the fixed and random effects is specified by the design matrices ***X*** and ***Z***, respectively.

For comparison, best linear unbiased estimates (BLUEs) for each volatile compound were also obtained, treating accession as a fixed rather than random effect:

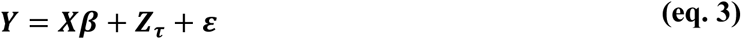

where ***τ*** corresponds to the vector of accession effects treated as fixed; for this model the solution is equivalent to the arithmetic mean of the replicates of each accession. All other terms are as defined in eq. 2. Both estimation approaches gave identical chemotype structure across all four quantification datasets (identical cluster membership, adjusted Rand index = 1.00; 100% overlap in significantly discriminating compounds; and the same optimal cluster number; Table S8). Values below the limit of detection were assigned LOD/2, defined per compound as half the minimum positive value observed for that compound. Imputed BLUP estimates were used for chemotype classification and machine learning, which require a complete data matrix. Descriptive statistics, compound detection frequencies and odor activity values were computed on the raw, unimputed data.

Differences among accessions were tested by one-way ANOVA with Bonferroni correction (*α* = 8.20×10⁻⁴). Effect sizes were reported as *η*² (Cohen, 2013), and significant pairwise differences were identified using Tukey’s HSD test. Because the number of technical replicates varied among accessions, unbalanced ANOVA was used. Pairwise Pearson correlation coefficients among compounds were visualized in heatmaps with hierarchical clustering (Ward Jr, 1963). Analysis of variance and correlation analysis were performed on imputed, log-transformed data.

#### 2.4.2 Chemotype classification

Prior to distance calculation and ordination, values below the limit of detection were assigned LOD/2 at the replicate level; BLUP estimates were then centred and scaled per compound (*z*-score) and clipped at ±3 SD. Compound-wise scaling equalises the contribution of compounds spanning different concentration ranges while retaining differences in total volatile output among accessions. The optimal number of clusters was determined using Calinski-Harabasz index (Caliński and Harabasz, 1974), elbow method (Ketchen and Shook, 1996), and silhouette coefficient (Rousseeuw, 1987). Hierarchical clustering analysis (HCA) was performed using Ward’s minimum variance method on a Euclidean distance matrix. Principal component analysis (PCA; Jolliffe and Cadima 2016) was used for dimensionality reduction and visualization; biplots displayed genotypes in PC1-PC2 space with color-coding according to HCA-derived clustering assignments (Jolliffe and Cadima, 2016). Clustering robustness was further assessed by t-distributed stochastic neighbor embedding (t-SNE; Maaten and Hinton 2008).

#### 2.4.3 Identification of discriminative compounds

Discriminatory volatiles were identified using three complementary feature selection methods: Mutual Information coupled with Support Vector Machine (MI-SVM) (Cortes and Vapnik, 1995; Kraskov et al., 2004), Recursive Feature Elimination-Support Vector Machine (RFE-SVM) (Guyon et al., 2002), and Random Forest importance ranking (Breiman, 2001). Consensus markers were defined as compounds selected by all three methods.

The dataset was split into training (80%) and testing (20%) set with stratified sampling. Classification performance was assessed using precision, recall (sensitivity), F1-score, specificity, balanced accuracy, Receiver Operating Characteristics Areas Under the Curve (ROC AUC) score (Fawcett, 2006) and confusion matrix, as defined in equations (4) – (8):

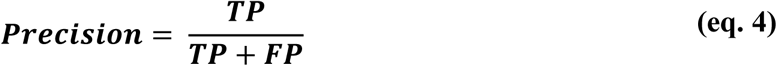

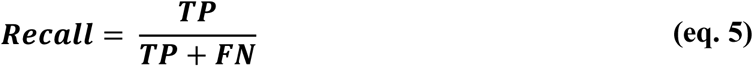

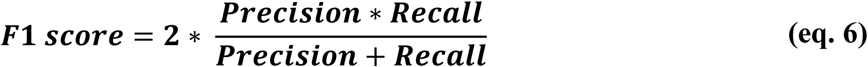

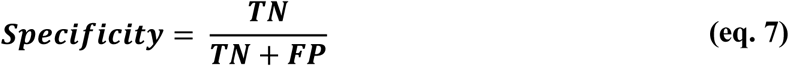

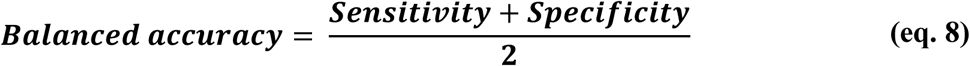

where:

*TP* (True Positives) = The number of accessions correctly classified as belonging to a specific chemotype cluster.
*FP* (False Positives) = The number of accessions incorrectly classified as belonging to a specific chemotype cluster.
*TN* (True Negatives) = The number of accessions correctly classified as not belonging to a specific chemotype cluster.
*FN* (False Negatives) = The number of accessions incorrectly classified as not belonging to a specific chemotype cluster when they belong to that cluster.

### 2.5 Software and visualizing

All data processing, statistical analyses, and visualizations were performed using R version 4.3.1 (Team, 2020) and Python version 3.12.4. BLUP and BLUE estimation were conducted with the *AllInOne* package (Najafabadi et al., 2023) in R. Python analyses utilized NumPy 1.24.3 (Harris et al., 2020) for numerical array operations; pandas 2.0.3 (McKinney, 2011) for data frame manipulation and preprocessing; SciPy 1.11.1 (Virtanen et al., 2020) for statistical tests, hierarchical clustering, and correlation analyses; scikit-learn 1.3.0 (Pedregosa et al., 2011) for machine learning algorithms including *k*-means clustering, hierarchical clustering, PCA, t-SNE, SVM, RF classification, Mutual Information, and Recursive Feature Elimination; statsmodels 0.14.0 (Seabold and Perktold, 2010) for advanced statistical modeling and post-hoc tests; Matplotlib 3.7.2 (Bisong, 2019) for base plotting functionality; seaborn 0.12.2 (Waskom, 2021) for statistical data visualization; and RDKit 2023.09.1 (Landrum, 2013) for cheminformatics analysis and molecular structure processing. Machine learning workflows followed the pipeline described by Babaei et al. (2026b), available at https://github.com/Mehdibabaeii/CannaFeatML.

## 3. RESULTS

### 3.1 Volatile organic compound (VOC) identification and chemical diversity

Gas chromatography analysis of 490 analytical runs from 165 drug-type cannabis accessions allowed the identified 61 high-confidence volatile organic compounds (VOCs) (Table 1, Fig. 1). Compounds were assigned to three main biosynthetic classes: terpenoids (*n* = 45; 73.8%), fatty acid-derived volatiles (*n* = 12; 19.7%), and amino acid-derived volatiles (*n* = 4; 6.6%) (Fig. S1a and Fig. 2a).

**Fig. 1.**
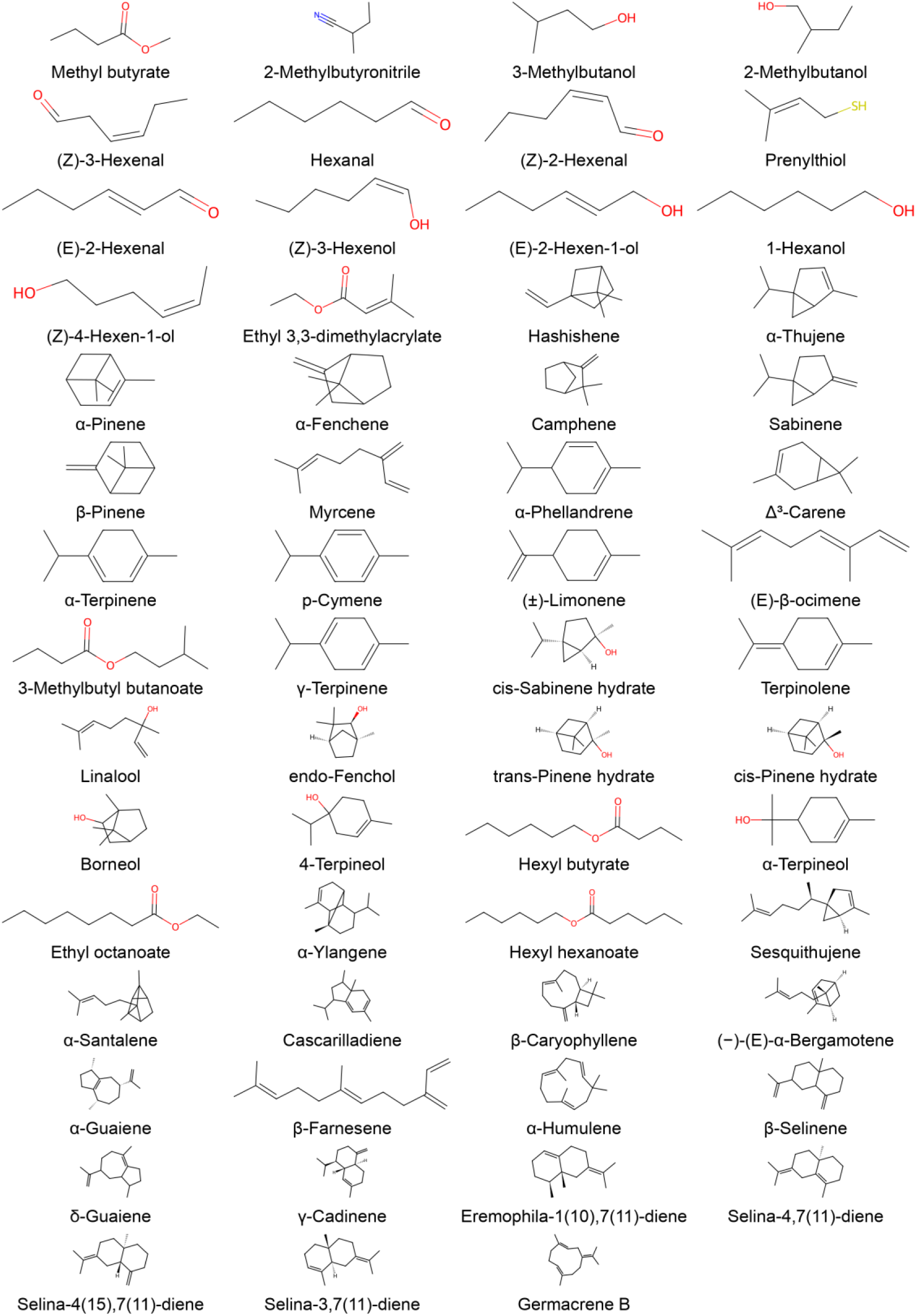
Chemical structures of volatile organic compounds (VOCs) identified in 165 cannabis accessions.

**Fig. 2.**
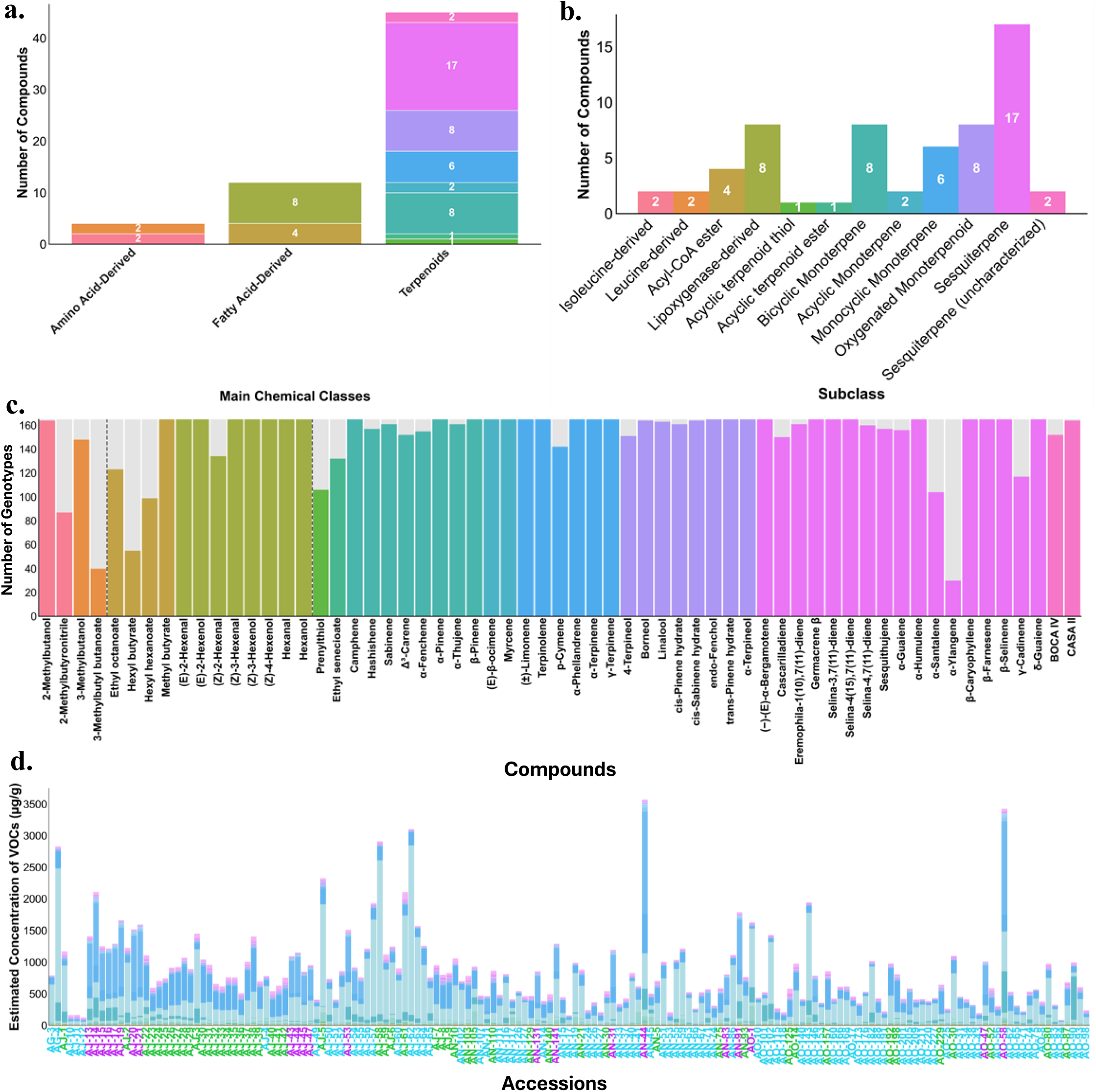
Chemical classification and distribution of volatile organic compounds (VOCs) in 165 drug-type cannabis accessions. (**a**–**b**) Number of VOCs across three main biosynthetic classes and 12 subgroups. (**c**) Detection frequency of 61 VOCs by subclass (% of accessions). (**d**) Estimated concentration profiles (µg/g fresh weight) across all accessions; accession labels are color-coded by chemotype (blue: Cluster ZERO, green: Cluster ONE, purple: Cluster TWO) as determined in Section 3.3.

**Table 1.** Volatile organic compounds (VOCs) identified in the drug-type cannabis accession.

| Peak No. | Retention time (min) | Compound | Main Class | Subgroup | Molecular Formula | Detection (%) | Mean ± SD ($\mu\text{g g}^{-1}$ FW) |
| --- | --- | --- | --- | --- | --- | --- | --- |
| 1 | 5.80 | Methyl butyrate | Fatty Acid-Derived | Acyl-CoA ester | C5H10O2 | 100 | 0.317 ± 0.162 |
| 2 | 5.92 | 2-Methylbutyronitrile | Amino Acid-Derived | Isoleucine-derived | C5H9N | 53 | 0.162 ± 0.046 |
| 3 | 6.02 | 3-Methylbutanol | Amino Acid-Derived | Leucine-derived | C5H12O | 90 | 0.195 ± 0.088 |
| 4 | 6.10 | 2-Methylbutanol | Amino Acid-Derived | Isoleucine-derived | C5H12O | 99 | 0.222 ± 0.109 |
| 5 | 7.44 | (Z)-3-Hexenal | Fatty Acid-Derived | Lipoxygenase-derived | C6H10O | 100 | 0.419 ± 0.431 |
| 6 | 7.48 | Hexanal | Fatty Acid-Derived | Lipoxygenase-derived | C6H12O | 100 | 1.629 ± 0.893 |
| 7 | 7.60 | (Z)-2-Hexenal | Fatty Acid-Derived | Lipoxygenase-derived | C6H10O | 81 | 0.172 ± 0.053 |
| 8 | 8.10 | Prenylthiol | Terpenoids | Acyclic terpenoid thiol | C5H10S | 64 | 0.145 ± 0.061 |
| 9 | 8.81 | (E)-2-Hexenal | Fatty Acid-Derived | Lipoxygenase-derived | C6H10O | 100 | 3.202 ± 2.317 |
| 10 | 8.89 | (Z)-3-Hexenol | Fatty Acid-Derived | Lipoxygenase-derived | C6H12O | 100 | 1.817 ± 1.122 |
| 11 | 9.15 | (E)-2-Hexenol | Fatty Acid-Derived | Lipoxygenase-derived | C6H12O | 100 | 2.925 ± 1.812 |
| 12 | 9.20 | Hexanol | Fatty Acid-Derived | Lipoxygenase-derived | C6H14O | 100 | 3.746 ± 2.198 |
| 13 | 9.26 | (Z)-4-Hexenol | Fatty Acid-Derived | Lipoxygenase-derived | C6H12O | 100 | 0.284 ± 0.163 |
| 14 | 10.70 | Ethyl senecioate | Terpenoids | Acyclic terpenoid ester | C7H12O2 | 80 | 1.737 ± 2.893 |
| 15 | 10.75 | Hashishene | Terpenoids | Bicyclic Monoterpene | C10H16 | 95 | 0.177 ± 0.061 |
| 16 | 10.93 | α-Thujene | Terpenoids | Bicyclic Monoterpene | C10H16 | 98 | 0.548 ± 0.991 |
| 17 | 11.20 | α-Pinene | Terpenoids | Bicyclic Monoterpene | C10H16 | 100 | 20.931 ± 35.682 |
| 18 | 11.53 | α-Fenchene | Terpenoids | Bicyclic Monoterpene | C10H16 | 94 | 0.182 ± 0.060 |
| 19 | 11.58 | Camphene | Terpenoids | Bicyclic Monoterpene | C10H16 | 100 | 2.019 ± 1.673 |
| 20 | 12.19 | Sabinene | Terpenoids | Bicyclic Monoterpene | C10H16 | 98 | 0.481 ± 0.871 |
| 21 | 12.34 | β-Pinene | Terpenoids | Bicyclic Monoterpene | C10H16 | 100 | 23.586 ± 20.380 |
| 22 | 12.68 | Myrcene | Terpenoids | Acyclic Monoterpene | C10H16 | 100 | 366.822 ± 458.080 |
| 23 | 13.03 | α-Phellandrene | Terpenoids | Monocyclic Monoterpene | C10H16 | 100 | 4.363 ± 9.898 |
| 24 | 13.22 | $\Delta^3$-Carene | Terpenoids | Bicyclic Monoterpene | C10H16 | 92 | 3.227 ± 7.895 |
| 25 | 13.33 | α-Terpinene | Terpenoids | Monocyclic Monoterpene | C10H16 | 100 | 3.026 ± 7.020 |
| 26 | 13.52 | p-Cymene | Terpenoids | Monocyclic Monoterpene | C10H14 | 86 | 0.280 ± 0.468 |
| 27 | 13.66 | (±)-Limonene | Terpenoids | Monocyclic Monoterpene | C10H16 | 100 | 171.814 ± 151.038 |
| 28 | 14.11 | (E)-β-ocimene | Terpenoids | Acyclic Monoterpene | C10H16 | 100 | 52.327 ± 101.001 |
| 29 | 14.36 | 3-Methylbutyl butanoate | Amino Acid-Derived | Leucine-derived | C9H18O2 | 24 | 0.269 ± 0.153 |
| 30 | 14.43 | γ-Terpinene | Terpenoids | Monocyclic Monoterpene | C10H16 | 100 | 1.502 ± 3.138 |
| 31 | 14.65 | cis-Sabinene hydrate | Terpenoids | Oxygenated Monoterpenoid | C10H18O | 99 | 0.352 ± 0.211 |
| 32 | 15.21 | Terpinolene | Terpenoids | Monocyclic Monoterpene | C10H16 | 100 | 109.903 ± 267.904 |
| 33 | 15.37 | Linalool | Terpenoids | Oxygenated Monoterpenoid | C10H18O | 99 | 19.185 ± 16.392 |
| 34 | 15.89 | endo-Fenchol | Terpenoids | Oxygenated Monoterpenoid | C10H18O | 100 | 4.204 ± 3.336 |
| 35 | 16.12 | trans-Pinene hydrate | Terpenoids | Oxygenated Monoterpenoid | C10H18O | 100 | 2.375 ± 1.831 |
| 36 | 16.53 | cis-Pinene hydrate | Terpenoids | Oxygenated Monoterpenoid | C10H18O | 98 | 0.426 ± 0.417 |
| 37 | 17.23 | Borneol | Terpenoids | Oxygenated Monoterpenoid | C10H18O | 99 | 0.623 ± 0.338 |
| 38 | 17.59 | 4-Terpineol | Terpenoids | Oxygenated Monoterpenoid | C10H18O | 92 | 0.544 ± 1.274 |
| 39 | 17.71 | Hexyl butyrate | Fatty Acid-Derived | Acyl-CoA ester | C10H20O2 | 33 | 1.105 ± 1.646 |
| 40 | 17.77 | α-Terpineol | Terpenoids | Oxygenated Monoterpenoid | C10H18O | 100 | 1.535 ± 0.999 |
| 41 | 18.02 | Ethyl octanoate | Fatty Acid-Derived | Acyl-CoA ester | C10H20O2 | 75 | 0.215 ± 0.315 |
| 42 | 21.95 | α-Ylangene | Terpenoids | Sesquiterpene | C15H24 | 18 | 0.514 ± 0.471 |
| 43 | 22.08 | Hexyl hexanoate | Fatty Acid-Derived | Acyl-CoA ester | C12H24O2 | 60 | 0.433 ± 0.639 |
| 44 | 22.70 | CASA II | Terpenoids | Sesquiterpene (uncharacterized) | - | 99 | 0.326 ± 0.183 |
| 45 | 22.98 | Sesquithujene | Terpenoids | Sesquiterpene | C15H24 | 95 | 0.276 ± 0.234 |
| 46 | 23.12 | α-Santalene | Terpenoids | Sesquiterpene | C15H24 | 63 | 0.326 ± 0.259 |
| 47 | 23.17 | Cascarilladiene | Terpenoids | Sesquiterpene | C15H24 | 91 | 0.364 ± 0.255 |
| 48 | 23.26 | β-Caryophyllene | Terpenoids | Sesquiterpene | C15H24 | 100 | 17.961 ± 15.960 |
| 49 | 23.41 | (-)-(E)-α-Bergamotene | Terpenoids | Sesquiterpene | C15H24 | 100 | 1.333 ± 1.248 |
| 50 | 23.54 | α-Guaiene | Terpenoids | Sesquiterpene | C15H24 | 95 | 0.848 ± 1.513 |
| 51 | 23.67 | β-Farnesene | Terpenoids | Sesquiterpene | C15H24 | 100 | 1.553 ± 1.836 |
| 52 | 23.83 | BOCA IV | Terpenoids | Sesquiterpene (uncharacterized) | - | 92 | 0.423 ± 0.923 |
| 53 | 23.98 | α-Humulene | Terpenoids | Sesquiterpene | C15H24 | 100 | 4.823 ± 3.992 |
| 54 | 24.66 | β-Selinene | Terpenoids | Sesquiterpene | C15H24 | 100 | 1.045 ± 0.776 |
| 55 | 24.98 | δ-Guaiene | Terpenoids | Sesquiterpene | C15H24 | 100 | 1.040 ± 1.791 |
| 56 | 25.14 | γ-Cadinene | Terpenoids | Sesquiterpene | C15H24 | 71 | $0.258 \pm 0.343$ |
| 57 | 25.36 | Eremophila-1(10),7(11)-diene | Terpenoids | Sesquiterpene | C15H24 | 98 | $0.469 \pm 0.312$ |
| 58 | 25.52 | Selina-4,7(11)-diene | Terpenoids | Sesquiterpene | C15H24 | 97 | $0.606 \pm 0.390$ |
| 59 | 25.63 | Selina-4(15),7(11)-<br>diene | Terpenoids | Sesquiterpene | C15H24 | 100 | $2.483 \pm 1.855$ |
| 60 | 25.78 | Selina-3,7(11)-diene | Terpenoids | Sesquiterpene | C15H24 | 100 | $2.405 \pm 1.974$ |
| 61 | 26.12 | Germacrene B | Terpenoids | Sesquiterpene | C15H24 | 100 | $2.804 \pm 2.577$ |

Within terpenoids, monoterpenes were the most numerous group (*n* = 24), comprising bicyclic (*n* = 8), oxygenated (*n* = 8), monocyclic (*n* = 6) and acyclic (*n* = 2) forms, followed by sesquiterpenes (*n* = 19, including two uncharacterized) and single representatives of acyclic terpenoid esters and acyclic terpenoid thiols (Fig. 2b). Fatty acid-derived volatiles were grouped in lipoxygenase-derived compounds (*n* = 8) and acyl-CoA esters (*n* = 4), while amino acid-derived volatiles were evenly divided between isoleucine-derived (*n* = 2) and leucine-derived (*n* = 2) compounds. Two sesquiterpenes (CASA II and BOCA IV) remained structurally uncharacterized despite consistent chromatographic detection.

Chemical prevalence varied substantially among accessions. Terpenoids were most consistently detected (93.6% of accessions), followed by fatty acid-derived volatiles (87.4%), whereas amino acid-derived volatiles showed the most restricted distribution (66.5%) (Fig. S1b). At the subclass level, acyclic monoterpenes were present in all accessions (100%) and oxygenated monoterpenoids in 98.3%, in contrast to more restricted distributions for acyl-CoA esters (67.0%), acyclic terpenoid thiols (64.2%) and leucine-derived compounds (57.0%) (Fig. S1c). Individual compound frequency ranged from ubiquitous detection (30 compounds detected in all 165 accessions, including myrcene, (±)-limonene, α-humulene and (Z)-3-hexenol) to sporadic occurrence (e.g., α- ylangene, *n* = 30; 3-methylbutyl butanoate, *n* = 40) (Fig. 2c). Comprehensive physicochemical descriptors, molecular properties, and database identifiers for all 61 compounds are provided in Table S6.

### 3.2 Accession-level variation in VOC profiles

Quantitative analysis of the 61 VOCs across 165 accessions revealed substantial accession-level variation (Table S9). Estimated concentrations (µg g⁻¹ FW) spanned more than three orders of magnitude, from low-abundance compounds (prenylthiol: 0.145 ± 0.061; 2-methylbutyronitrile: 0.162 ± 0.046) to dominant monoterpenes (myrcene: 366.82 ± 458.08; (±)-limonene: 171.81 ± 151.04). At the class level, mean abundance was highest for terpenoids (19.71 ± 106.71), followed by fatty acid-derived volatiles (1.46 ± 1.81) and amino acid-derived volatiles (0.21 ± 0.10). Within terpenoids, acyclic monoterpenes exhibited the highest subclass abundance (209.58 ± 366.72), primarily driven by the dominance of myrcene, exceeding monocyclic monoterpenes (49.63 ± 143.98), bicyclic monoterpenes (6.55 ± 17.65), oxygenated monoterpenoids (3.68 ± 8.48) and sesquiterpenes (2.51 ± 6.12).

Stacked percentage compositions visualized the proportional contribution of individual VOCs to each accession’s profile (Fig. 2d). Abundance distributions exhibited extensive variability, with coefficients of variation ranging from 0.29 to 2.45 for individual compounds and 0.42 to 2.90 at the subclass level. Interquartile ranges varied from tightly constrained (prenylthiol: IQR = 0.015; 2-methylbutyronitrile: IQR = 0.045) to highly dispersed (myrcene: IQR = 309.02; (±)-limonene: IQR = 180.53), reflecting substantial accession-level differences in VOC accumulation (Table S9).

One-way ANOVA confirmed highly significant accession effects for all 61 compounds (*p* < 0.001, Table S10), with effect sizes (*η*²) ranging from 0.67 to 0.97, indicating that between-accession differences accounted for 67–97% of the total measured variation. Repeatability estimates, calculated as the proportion of total variance attributable to accession, were moderate to high (mean = 0.81, range: 0.50–0.95; Table S11). The highest values were observed for hexyl hexanoate (*R* = 0.95, *η*² = 0.97), (E)-β-ocimene (*R* = 0.94, *η*² = 0.96), Δ³-carene and 3-methylbutyl butanoate (both *R* = 0.93). In contrast, several C6 aldehydes and alcohols showed lower repeatability ((E)-2-hexenol: *R* = 0.50, *η*² = 0.67; hexanal: *R* = 0.55, *η*² = 0.70), consistent with their higher analytical variability among technical replicates.

### 3.3 Chemotype classification

#### 3.3.1 Unsupervised clustering and multivariate characterization

Cluster validation metrics gave partially divergent results: the Calinski-Harabasz index reached its maximum at *k* = 3 (26.31) and the elbow method indicated an inflection at *k* = 3, whereas the silhouette coefficient was marginally higher for *k* = 2 (0.203) than for *k* = 3 (0.2005). Both silhouette values are low, indicating that the partition is not sharply separated. A three-cluster solution was adopted because the elbow criterion gave *k* = 3 consistently across all four quantification datasets, cluster membership was identical under BLUP and accession-mean estimation (Table S8), and an independent supervised classifier trained on a subset of compounds recovered the three groups with 97% accuracy on held-out accessions (Section 3.3.2). Unsupervised hierarchical clustering of volatile profiles partitioned the 165 accessions into three chemotypes: Cluster ZERO (*n* = 90, 54.5%), Cluster ONE (*n* = 53, 32.1%), and Cluster TWO (*n* = 22, 13.3%) (Table S12). Hierarchical cluster analysis using Euclidean distance and Ward’s linkage revealed clear separation among the three groups (Fig. 3a), corroborated by principal component analysis where PC1 and PC2 explained 39.1% of total variance (22.5% and 16.6%, respectively; Fig. S2a). t-distributed stochastic neighbor embedding (t-SNE) further confirmed distinct clustering patterns with minimal overlap among chemotypes (Fig. S2b).

**Fig. 3.**
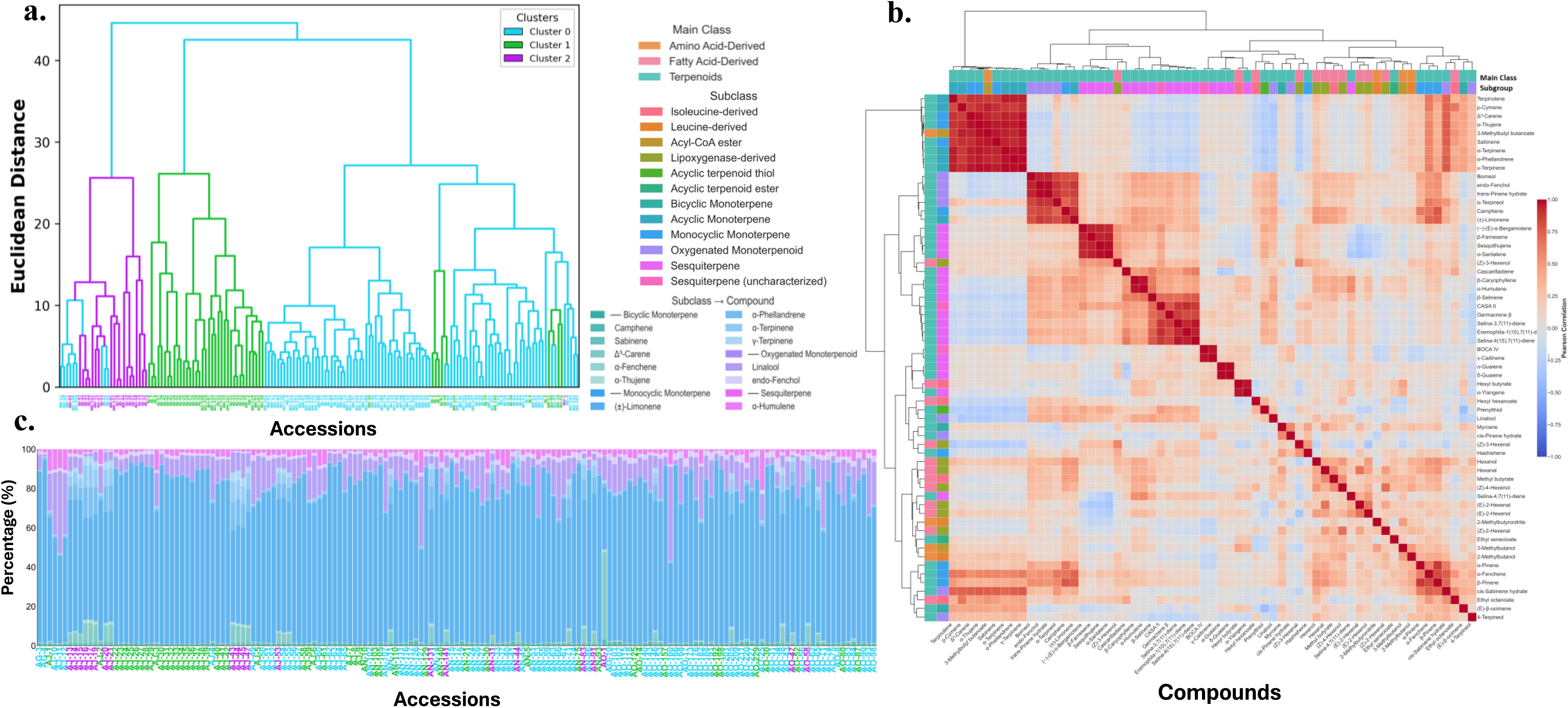
Chemotype classification of 165 drug-type cannabis accessions. (**a**) Hierarchical clustering dendrogram based on 61 VOC profiles revealing three data-derived chemotypes (Cluster ZERO, *n* = 90; Cluster ONE, *n* = 53; Cluster TWO, *n* = 22). (**b**) Pearson correlation matrix of 61 VOCs within and across biosynthetic classes. (**c**) Accession distribution across three chemotypes based on 12 discriminative compounds (Camphene, α-Fenchene, Sabinene, α-Terpinene, (±)-Limonene, α-Humulene, Linalool, endo-Fenchol, Δ³-Carene, α- Thujene, γ-Terpinene, and α-Phellandrene) identified through machine learning-based feature selection.

Chemotype-specific profiles revealed marked differences in volatile signatures (Fig. S2c, Table S13). Cluster ZERO was characterized by consistently low concentrations across most compounds, including (±)-limonene (90.79 vs. 314.37 in Cluster ONE; *F* = 266.62, *d* = −1.43, *p* < 0.001), α- fenchene (*F* = 299.19, *d* = −1.53) and camphene (*F* = 200.04, *d* = −1.24). Cluster ONE was enriched in camphene (3.61 vs. 1.21 in Cluster ZERO; *F* = 373.79, *d* = +1.65), (±)-limonene (*F* = 362.00, *d* = +1.64), β-caryophyllene (32.12 vs. 11.01 in Cluster ZERO; *F* = 297.80, *d* = +1.44), α-humulene (*F* = 281.80, *d* = +1.42); linalool was markedly enriched in Cluster ONE (32.4), exceeding both Cluster ZERO (13.09) and Cluster TWO (10.37; *F* = 250.13, *d* = +1.33). Cluster TWO, the smallest group, was distinguished by exceptionally high terpinolene (668.18, >20-fold higher than the other chemotypes; *F* = 1013.47, *d* = +2.29) together with elevated minor monoterpenes (α-thujene: *F* = 1062.96, *d* = +2.37; α-terpinene: *F* = 892.96, *d* = +2.13; α-phellandrene: *F* = 888.93, *d* = +2.13; γ- terpinene: *F* = 873.98, *d* = +2.11; Δ³-carene: *F* = 834.25, *d* = +2.03) and 3-methylbutyl butanoate (*F* = 1220.93, *d* = +2.65), all *p* < 0.001.

Correlation analysis of the 61 VOCs revealed complex co-regulation patterns within and across chemical classes (Fig. 3b). Strong positive correlations (*r* = 0.95–0.99) were observed among co- biosynthesized terpenoids, including oxygenated monoterpenoids (endo-fenchol vs. trans-pinene hydrate: *r* = 0.992), monocyclic monoterpenes (α-terpinene vs. γ-terpinene: *r* = 0.991; α- phellandrene vs. γ-terpinene: *r* = 0.977) and sesquiterpenes (α-guaiene vs. δ-guaiene: *r* = 0.990; sesquithujene vs. α-santalene: *r* = 0.981; β-caryophyllene vs. α-humulene: *r* = 0.976). Correlations between monoterpene and sesquiterpene subclasses were generally weak, with occasional negative associations (Germacrene B vs. p-Cymene: *r* = −0.12). Notably, 3-methylbutyl butanoate, an amino acid-derived volatile, showed unexpectedly strong cross-class correlations with monoterpenes (*r* = 0.88–0.98), particularly with α-thujene (*r* = 0.977), γ-terpinene (*r* = 0.971), sabinene (*r* = 0.964), α-terpinene (*r* = 0.953) and Δ³-carene (*r* = 0.943), alongside hexyl butyrate with the sesquiterpene α-ylangene (*r* = 0.962). Hierarchical clustering of the accession-by-compound matrix was consistent with the three-chemotype assignments (Fig. S3), with Cluster TWO showing the most pronounced positive deviations (mean *Z* = +0.35, driven by Δ³-carene, α-thujene and γ-terpinene, all mean *Z* > 2.19), Cluster ONE showing moderate enrichment (mean *Z* = +0.29, led by camphene, β-caryophyllene and α-humulene), and Cluster ZERO representing the baseline profile (mean *Z* = −0.28).

#### 3.3.2 Supervised feature selection

Three supervised feature-selection algorithms were applied to identify minimal discriminative markers: Mutual Information coupled with Support Vector Machine (MI-SVM; 16 compounds; Table S14), Recursive Feature Elimination with SVM (RFE-SVM; 16 compounds; Table S15), and Random Forest (RF; 45 compounds ranked (12 trees); Table S16). Pairwise overlap ranged from 12 to 16 compounds, and the three-way intersection yielded 12 consensus markers: camphene, α- fenchene, sabinene, α-terpinene, (±)-limonene, α-humulene, linalool, endo-fenchol, Δ³-carene, α- thujene, γ-terpinene and α-phellandrene (Table S17).

On the held-out partition (*n* = 33), the 12-compound panel reproduced the full-profile chemotype assignment for 32 of 33 accessions (Cluster ONE: 10/11; Cluster TWO: 4/4; Cluster ZERO: 18/18), matching the MI-SVM and RFE-SVM feature sets (both 32/33) and exceeding the Random Forest panel (30/33). Because the chemotype labels were derived from the same measurements, this quantifies how compactly the 61-compound structure can be represented by a reduced panel rather than estimating prospective authentication accuracy; feature selection was performed before partitioning, so the estimate is optimistic. Single-marker discrimination was assessed independently by five-fold stratified cross-validation (Fig. S4a, S4b). Single-marker ROC analyses revealed chemotype-specific discrimination (Fig. S4a). Cluster TWO was identified with perfect accuracy by Δ³-carene (AUC = 1.000), α-thujene (0.998) and sabinene (0.979), while Cluster ONE was best discriminated by camphene (0.905), (±)-limonene (0.905) and α-humulene (0.887). Several terpinolene-type markers showed AUC values below 0.5 for Cluster ONE, reflecting discrimination by depletion rather than enrichment. Mean ROC curves identified α- fenchene (0.856) and linalool (0.766) as the strongest overall discriminators (Fig. S4b).

Compositional profiles of the 12 consensus compounds revealed distinct chemotype signatures (Fig. S4c), with clear accession-level separation (Fig. 3c) and tight intra-chemotype variance (Fig. S4d). These findings indicate that a parsimonious 12-compound panel captures the chemotype structure defined by the full profile, a prerequisite for developing targeted screening tools once validated on independent material.

### 3.4 Odor profiling and aromatic characterization

Odor activity values (OAVs) were calculated as the ratio of estimated concentration to literature-derived odor detection thresholds (ODTs) for the 33 compounds with available threshold data (Table S7). OAVs spanned more than six orders of magnitude, from camphene (0.078) to prenylthiol (284,159) (Fig. 4a, Fig. S5). The top-ranking odorants were prenylthiol (284,159), α- pinene (58,143), (±)-limonene (14,318), linalool (13,626) and myrcene (8,947), representing sulfur, pine, citrus, floral and herbal sensory categories, respectively. Scatter plot analysis of concentration versus OAV revealed distinct odorant groups: compounds with both high concentration and high OAV (e.g., myrcene, (±)-limonene, α-pinene) occupied the upper-right region, whereas prenylthiol emerged as a low-concentration compound with exceptionally high OAV, ranking far above more abundant compounds on this index (Fig. 4a). Among sensory descriptors, fruity (*n* = 7) and green (*n* = 4) were the most frequent (Fig. 4b).

**Fig. 4.**
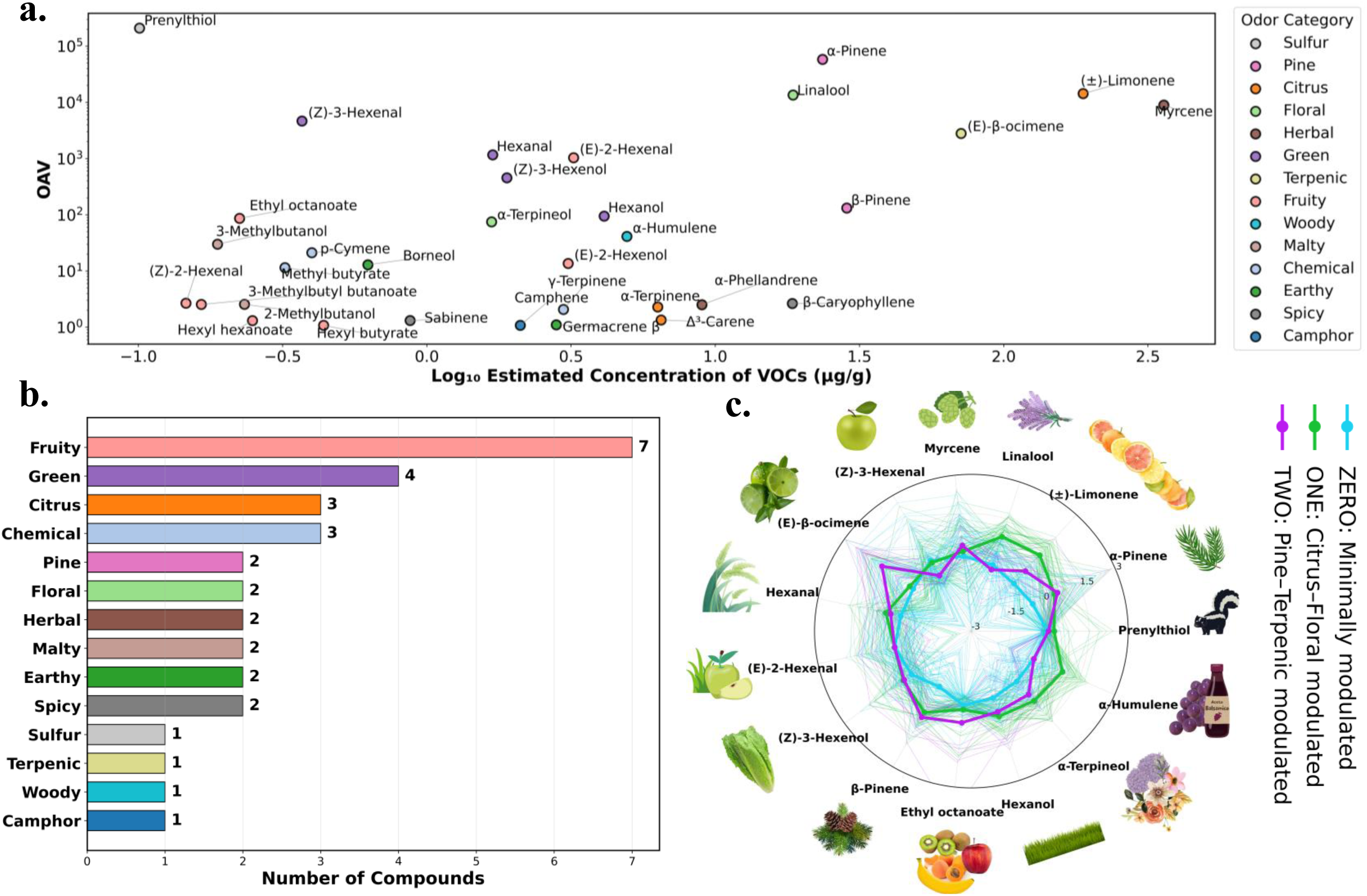
Odor activity value (OAV) analysis of 165 drug-type cannabis accessions. (**a**) Estimated concentration (log₁₀, µg/g) versus OAV for 33 odor-active compounds, color-coded by 14 sensory descriptors. (**b**) Number of compounds per descriptor category. (**c**) Radar plot of the 15 most odor-active compounds (Pareto-scaled log OAV); thin lines are individual accessions where detected, thick lines chemotype means (ZERO, *n* = 90, minimally modulated; ONE, *n* = 53, citrus–floral modulated; TWO, *n* = 22, pine–terpenic modulated).

Significant inter-chemotype differences were observed for 27 of the 33 compounds (2 at *p* < 0.05, 2 at *p* < 0.01 and 23 at *p* < 0.001; Kruskal–Wallis, Table S18, Fig. S5). All three chemotypes shared a prenylthiol-dominated core accounting for 67–77% of summed OAV and were therefore distinguished by the extent and nature of terpenoid modulation of that core, quantified as the non-sulfur fraction of summed OAV: Cluster ZERO (23.5%), Cluster ONE (29.6%) and Cluster TWO (32.8%). On this basis the chemotypes were designated *minimally modulated* (ZERO), *citrus– floral modulated* (ONE) and *pine–terpenic modulated* (TWO).

One-versus-rest testing identified the compounds characteristic of each chemotype (Table S18). Cluster ONE was enriched in (±)-limonene (26,902 vs. 7,164 in Cluster ZERO; *δ* = +0.83) and linalool (24,277 vs. 8,891; *δ* = +0.76), consistent with its citrus–floral designation. Cluster TWO was enriched in (E)-β-ocimene (7,497 vs. 2,318; *δ* = +0.56) and α-pinene (80,852 vs. 44,606; *δ* = +0.40) and showed the highest pine and terpenic category contributions. Cluster ZERO showed no significantly elevated odorant and instead exhibited the lowest values across most compounds, reflecting minimal terpenoid modulation. Notably, prenylthiol OAV did not differ significantly among chemotypes (Kruskal–Wallis, *ns*), confirming that these designations describe compositional balance rather than absolute odor intensity (Fig. 4c, Fig. S6).

## 4. DISCUSSION

This study presents a comprehensive volatilomics characterization of 165 drug-type *Cannabis sativa* accessions, identifying and quantifying 61 VOCs and establishing a robust three-chemotype classification framework grounded in quantitative profiling. Terpenoids dominated the identified compounds (73.8%), with monoterpenes (*n* = 24) and sesquiterpenes (*n* = 19) as the primary subgroups. This composition aligns with the biochemical specialization of cannabis glandular trichomes, where secondary metabolite production occurs via the plastidial methylerythritol phosphate (MEP) and cytosolic mevalonate (MVA) pathways, both generating C_5_ precursors (IPP and DMAPP) that condense to form geranyl diphosphate (GPP, C_10_) for monoterpenes and farnesyl diphosphate (FPP, C_15_) for sesquiterpenes (Ashour et al., 2010; Semmar, 2024; Wink, 2010). Comparable terpenoid dominance has been observed in smaller panels, such as 19 cultivars analyzed by HS-SPME-GC-MS (Janta and Vimolmangkang, 2024). The cannabis genome encodes 55 characterized TPS genes with tissue-specific expression patterns, with over 100 distinct terpenes identified to date and biosynthesis governed by at least 30 functional genes (Allen et al., 2019; Kaminski et al., 2025).

In the present study, unsupervised hierarchical clustering partitioned the accessions into three distinct chemotypes (*n* = 90, 53, and 22), with principal component analysis confirming clear separation (PC1 + PC2 explaining 39.1% of variance). This variance is consistent with prior reports on larger panels, where increased genetic diversity reduces explained variance (e.g., 36.8% across 460 accessions; Hazekamp et al. 2016), and contrasts with higher values in smaller panels (e.g., 61.2% across 19 cultivars yielding five clusters; Janta and Vimolmangkang 2024). Volatile profiles provided superior discriminating power for chemotype classification compared to cannabinoid ratios, as demonstrated by Hazekamp et al. (2016), who found that “Sativa” and “Indica” types showed comparable THC and CBD content but exhibited significant differences in terpene composition, particularly in hydroxylated sesquiterpenes (guaiol, beta-eudesmol, gamma- eudesmol) that were strongly associated with “Indica” types. β-Caryophyllene, the most abundant sesquiterpene in the present panel, showed the strongest enrichment in Cluster ONE (*d* = +1.44), consistent with Janta and Vimolmangkang (2024), who reported it as the predominant sesquiterpene in 12 of 19 cultivars (12.89–34.72% of total volatiles), and Russo (2011), who described it as the most common terpenoid in cannabis extracts. This pattern also aligns with Oswald et al. (2023), where the analysis of 31 ice-hash rosin extracts sourced from commercial dispensaries in Los Angeles, California, identified β-myrcene, D-(+)-limonene, β-caryophyllene, and terpinolene as the most abundant volatile constituents across varieties, reinforcing β- caryophyllene’s position among the major high-impact terpenes in cannabis. The chemotypes also differed in total volatile output (median 462, 858 and 1,292 µg g⁻¹ for Clusters ZERO, ONE and TWO), indicating that the classification reflects overall volatile productivity as well as compositional pattern.

Machine learning-based feature-selection identified a parsimonious set of 12 volatile markers consistently selected across three independent algorithms (MI, RFE, and RF importance): camphene, α-fenchene, sabinene, α-terpinene, (±)-limonene, α-humulene, linalool, endo-fenchol, Δ³-carene, α-thujene, γ-terpinene, and α-phellandrene. This consensus panel recovered the chemotype assignment of 32 of 33 held-out accessions, matching the performance of the full MI-SVM and RFE-SVM feature sets. In comparison, Janta and Vimolmangkang (2024) identified 20 discriminant markers via PLS-DA (VIP >1.0) (e.g., eucalyptol, (+)-2-carene, o-cymene, terpinolene, γ-eudesmol, α-bisabolol, humulene) in a 19-cultivar panel. The selected volatile markers are predominantly monoterpenes (C₁₀), spanning bicyclic (camphene, α-fenchene, sabinene, Δ³-carene, α-thujene), monocyclic ((±)-limonene, α-terpinene, γ-terpinene, α- phellandrene) and oxygenated (linalool, endo-fenchol) forms, together with the sesquiterpene α- humulene, arising from TPS-catalyzed cyclization cascades (Booth et al., 2017). Some marker pairs were negatively correlated, consistent with metabolic competition for shared precursors (GPP, FPP) or reciprocal transcriptional regulation, as demonstrated for cannabinoid biosynthesis where THCA and CBDA production exhibit competitive flux partitioning from their common precursor CBGA (Babaei and Torkamaneh, 2026a; Hesami et al., 2020; Kaminski et al., 2025). These findings are subject to the limits of the present design: the held-out partition contained only four Cluster TWO accessions, and replicates were technical rather than biological, so validation on external cohorts and repeated harvests will be required.

Odor activity value (OAV) analysis provides a quantitative framework bridging analytical chemistry and sensory perception (Patton and Josephson, 1957; Ruth, 1986; Wu et al., 2015). OAV application to cannabis using simultaneous multidimensional GC-MS-olfactometry demonstrated that abundant volatiles are not necessarily the most sensorially impactful—a paradigm shift from concentration-based assumptions (Rice and Koziel, 2015). Prior study revealed that compounds present at low absolute concentrations—including benzaldehyde, nonanal, and o-cymene— possessed higher OAV values than major constituents like β-myrcene and limonene, indicating disproportionate contribution to overall aroma despite minimal chemical abundance (Rice and Koziel, 2015). Similarly, in the present study several low-abundance compounds exhibited substantially elevated OAV values (e.g., Prenylthiol), confirming that relative odor potential is not predicted by concentration alone. Since concentrations are estimated from headspace-collected volatiles and detector response varies with compound structure, OAV values indicate relative rather than absolute sensory potential.

Among the 12 consensus markers, only nine had published odour detection thresholds, and their OAV ranks were markedly polarised: (±)-limonene (#3) and linalool (#4) ranked among the most odour-active compounds, whereas six others—α-phellandrene (#25), α-terpinene (#26), γ- terpinene (#27), Δ³-carene (#29), sabinene (#30) and camphene (#33)—ranked in the lowest third, with OAV values below 1.5. This dissociation indicates that chemotype classification integrates two distinct categories of marker: a small number of high-impact aroma compounds and a larger set of low-odour discriminatory markers that nonetheless discriminate accessions reliably (Fischedick, 2017; Janta and Vimolmangkang, 2024; Jin et al., 2021). This pattern may reflect historical selection of distinctive organoleptic profiles while maintaining chemotaxonomic consistency (Clarke and Merlin, 2016; Vergara et al., 2016).

The volatilomics framework established here addresses fundamental limitations in cannabis product standardization and breeding efficiency. Vernacular strain nomenclature exhibits pervasive unreliability, with identical names applied to genetically and chemically distinct materials—a problem undermining quality control, consumer protection, and regulatory compliance across legal markets (Schwabe and McGlaughlin, 2019). The chemotype classification system developed here, grounded in 61 quantified volatiles and recovery of the chemotype structure by a 12-compound panel, provides an objective categorization that could complement strain names. This approach parallels metabolomics-driven classification frameworks in related aromatic crops, particularly hops (*Humulus lupulus* L.), where volatile profiling enables cultivar authentication and hops chemistry prediction independent of environmental variation (Steenackers et al., 2015). The 12- compound consensus marker panel identified through integrated FS could support streamlined chemotype screening after independent validation (Fischedick, 2017; Janta and Vimolmangkang, 2024). Beyond authentication, these volatile traits could support phenotypic selection in breeding programs—a strategy proven effective across crop species for accelerating genetic gain while reducing phenotyping costs (Belzile et al., 2020; Torkamaneh et al., 2018). Integration of these phenotypic markers with emerging genomic tools, including QTL mapping for terpene biosynthesis loci (Babaei and Torkamaneh, 2026b; de Ronne and Torkamaneh, 2025) and genomic selection models (Yoosefzadeh Najafabadi and Torkamaneh, 2025) could enable marker-assisted selection once genotype–phenotype associations are established, and expedite cultivar development tailored to specific market segments defined by organoleptic preferences. However, restricted terpene diversity in contemporary commercial germplasm, attributed to intensive clandestine breeding and prohibition-era bottlenecks (Babaei et al., 2025; Clarke and Merlin, 2016; Kaminski et al., 2025), necessitates systematic germplasm expansion through evaluation of underutilized landrace accessions and hemp varieties (Babaei et al., 2024; Babaei and Ajdanian, 2020). Future research priorities include transcriptomic profiling to elucidate TPS gene expression dynamics underlying chemotype differentiation (Allen et al., 2019; Kaminski et al., 2025), metabolic flux analysis using stable isotope tracers to quantify carbon partitioning between terpene and cannabinoid biosynthesis (Bacher et al., 2016; Booth et al., 2017), genome-wide association studies (GWAS) to identify genomic loci controlling volatile production and enable precision breeding, and validation of chemotype perceptual relevance through trained sensory panels and consumer preference studies (Babaei and Torkamaneh, 2026a; Belzile and Torkamaneh, 2022; Torkamaneh and Belzile, 2022). These efforts will collectively enhance the genetic foundation, breeding precision, and market differentiation of cannabis cultivars while establishing reproducible quality standards essential for pharmaceutical and nutraceutical applications.

## 5. CONCLUSION

This comprehensive volatilomics study characterises the volatile diversity of 165 drug-type cannabis accessions, resolving it into three discrete chemotypes efficiently captured by a parsimonious 12-compound consensus marker set. The integration of analytical chemistry (GC-FID/MS with 61 quantified volatiles), multivariate classification, consensus feature selection, and estimated odor-activity screening provide a quantitative description of volatile diversity in this panel. Consumer preference, sensory perception, regulatory classification and reproducibility across environments were not assessed here. Establishing whether these chemotypes are stable across harvests and growing conditions, and whether they correspond to perceptible aroma differences, would be required before the framework could inform breeding decisions, product standardisation or authentication.

## Data Statement

Replicate-level volatile compound data for all four quantification approaches are available via Figshare (DOI: <u>10.6084/m9.figshare.33359469</u>). Processed data supporting the findings of this study are provided in the Supplementary Tables.

## CRediT authorship contribution statement

**Mehdi Babaei**: Conceptualization, Methodology, Investigation, Data curation, Formal analysis, Visualization, Writing – original draft, Writing – review and editing. **Charles Goulet**: Methodology, Resources, Supervision, Writing – review and editing. **Davoud Torkamaneh**: Conceptualization, Funding acquisition, Project administration, Resources, Supervision, Writing – review and editing.

## Acknowledgements

The authors wish to thank Alexis St-Gelais and Hubert Marceau (PhytoChemia, Chicoutimi, QC, Canada) for chromatographic peak integration, and Justine Richard-Giroux and Éliana Lapierre for plant growth, cultivation and sampling.

## Abbreviations

BLUE: Best Linear Unbiased Estimation
BLUP: Best Linear Unbiased Prediction
FW: Fresh Weight
GC-FID: Gas Chromatography-Flame Ionization Detection
GC-MS: Gas Chromatography-Mass Spectrometry
HCA: Hierarchical Clustering Analysis
HSD: Honestly Significant Difference (Tukey’s)
ISTD: Internal Standard
LOD: Limit of Detection
MI-SVM: Mutual Information-Support Vector Machine
ML: Machine Learning
OAV: Odor Activity Value
ODT: Odor Detection Threshold
PCA: Principal Component Analysis
RF: Random Forest
RFE-SVM: Recursive Feature Elimination-Support Vector Machine
ROC-AUC: Receiver Operating Characteristic-Area Under the Curve
t-SNE: t-Distributed Stochastic Neighbor Embedding
VOC: Volatile Organic Compound.

