## Supplemental Figures S1 to S6 for "Volatile profiling and estimated odor-activity analysis of commercial drug-type cannabis accessions"

**Fig. S1** Chemical classification of 61 volatile organic compounds (VOCs) identified across 165 drug-type cannabis accessions. (**a**) Relative proportion of three main biosynthetic classes (inner) and 12 subgroups (outer). (**b**) Detection frequency of main classes expressed as percentage of accessions. (**c**) Detection frequency of 12 biosynthetic subgroups expressed as percentage of accessions.

**Fig. S2** Chemotype classification and VOC profiles of 165 drug-type cannabis accessions based on 61 VOCs. **(a)** Principal component analysis (PCA) biplot showing separation of three chemotypes (Cluster ZERO, *n* = 90; Cluster ONE, *n* = 53; Cluster TWO, *n* = 22). **(b)** t-distributed stochastic neighbor embedding (t-SNE) visualization of chemotype clustering. **(c)** Mean estimated concentration profiles (µg/g fresh weight) of 61 VOCs across three chemotypes, grouped by biosynthetic class and subclass.

**Fig. S3** Hierarchical clustering heatmap of 165 drug-type cannabis accessions × 61 volatile compounds, showing distinct compositional signatures of the three chemotypes (Cluster ZERO, *n* = 90; Cluster ONE, *n* = 53; Cluster TWO, *n* = 22). Values are *Z*-scores computed per compound across all accessions.

**Fig. S4** Machine learning-based chemotype classification of 165 drug-type cannabis accessions using 12 discriminative VOCs. **(a)** Individual chemotype ROC curves for each of the three chemotypes. **(b)** Mean ROC curves of 12 consensus compounds selected by MI-SVM, RFE-SVM, and RF methods. **(c)** Mean estimated concentration profiles (µg/g fresh weight) of 12 discriminative VOCs across three chemotypes (Cluster ZERO, *n* = 90; Cluster ONE, *n* = 53; Cluster TWO, *n* = 22). **(d)** Box plot distributions of 12 key VOCs (Camphene, α-Fenchene, Sabinene, α-Terpinene, (±)-Limonene, α-Humulene, Linalool, endo-Fenchol, Δ³-Carene, α-Thujene, γ-Terpinene, and α-Phellandrene) differentiating cannabis chemotypes.

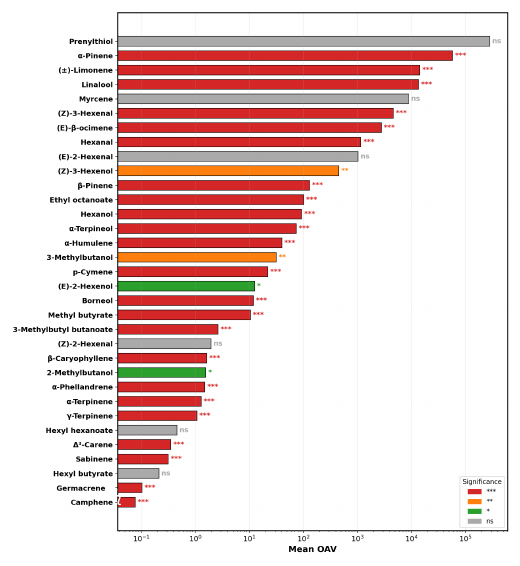

**Fig. S5** Mean odor activity values (OAV) of 33 odor-active compounds across 165 drug-type cannabis accessions, ranked in descending order. Bar colors indicate significance of differences among the three chemotypes (Kruskal-Wallis; ns: not significant, **p* < 0.05, ***p* < 0.01, ****p* < 0.001).

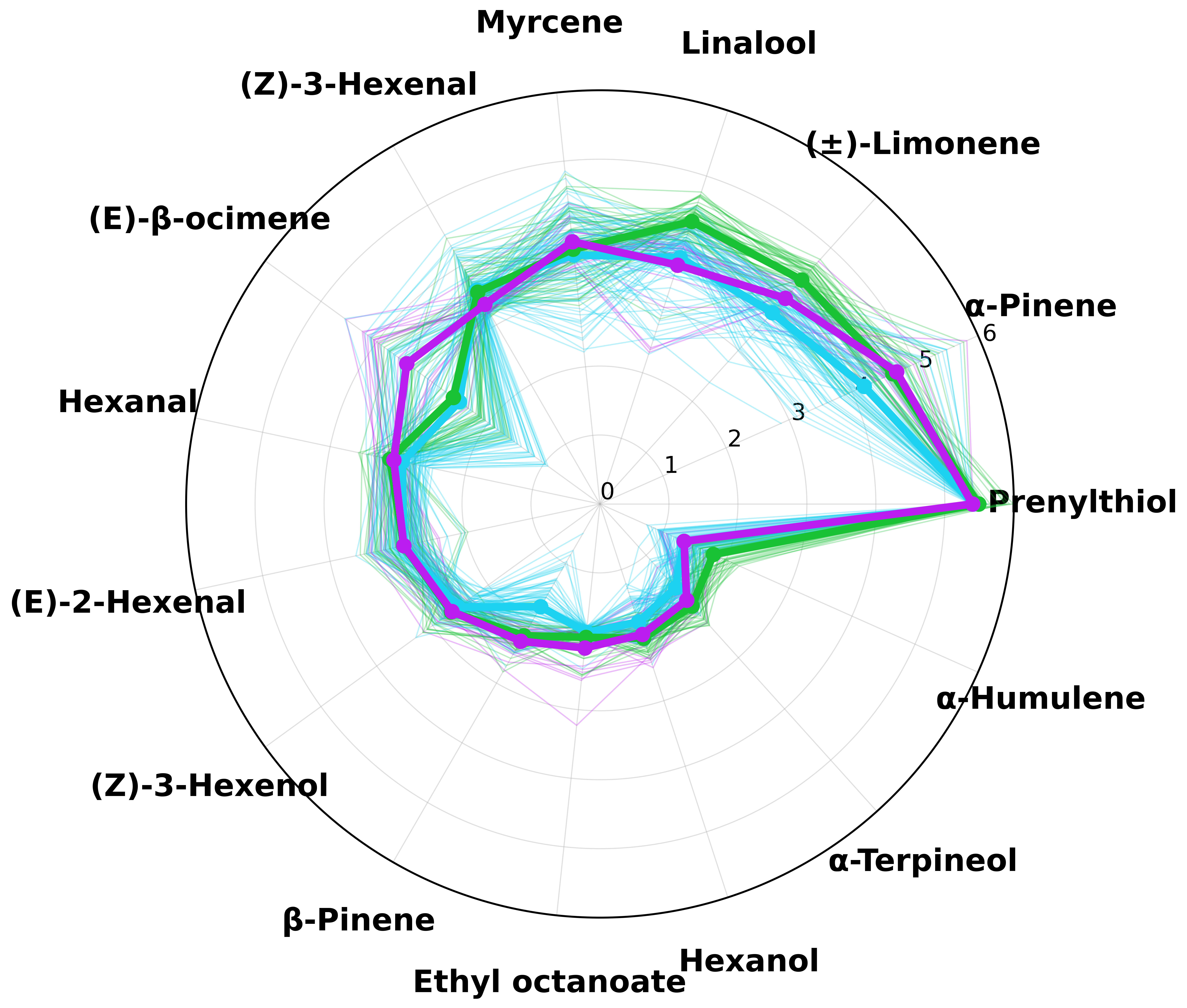

**Fig. S6** Chemotype odor-activity profiles of the 15 most odor-active compounds, shown as log₁₀(OAV) without scaling. Thin lines are individual accessions colored by chemotype; thick lines are chemotype means (ZERO, *n* = 90; ONE, *n* = 53; TWO, *n* = 22). All three chemotypes share a prenylthiol-dominated core, indicating that the separation in Fig. 4c reflects compositional balance rather than absolute odor intensity.
